# Increasing environmental fluctuations can dampen variability of endogenously cycling populations

**DOI:** 10.1101/2023.05.10.531506

**Authors:** Nicholas Kortessis, José Miguel Ponciano, Franz W. Simon, Jake M. Ferguson

## Abstract

Understanding how populations respond to increasingly variable conditions is a major objective for natural resource managers attempting to forecast extinction risk. The lesson from current modeling is clear: Increasing environmental variability increases population abundance variability. We show that this paradigm fails to describe a broad class of empirically observed dynamics, namely endogenously-driven population cycles. In contrast to the dominant paradigm, these populations can exhibit reduced long-run population variance under increasing environmental variability. We hypothesize that this paradox arises from interactions between environmental stochasticity and nonlinear density dependence. Such interactions violate the oft-assumed additivity of stochastic and deterministic drivers of population fluctuations present in many models that forecast population size. We show evidence for the interaction in two canonical cyclical populations: flour beetles and Canadian Lynx. To help identify the interaction, we develop new theory to quantify the strength of these interactions by partitioning the effects of nonlinear dynamics and stochastic variation on dynamical systems. In both empirical examples, the partitioning shows that the interaction between deterministic and stochastic dynamics reduces the overall variance in population size. Our results highlight that previous predictions about extinction under environmental variability may prove inadequate to understand the effects of climate change in many populations.

## Introduction

Most projections of future biodiversity under climate change present a bleak outlook [1,2, 3,4,5]. Models predict that the warmer average temperatures predicted by climate models will be outside the current thermal limits of some species [6,7]. While changes in average conditions are likely to have significant consequences for many species, more frequent extreme events are also predicted to occur with climatic change. Examples include the projected increases in variability of annual temperature [8] and rainfall [9]. In a landmark study, Vasseur et al. [10] used thermal performance curves to show that future changes in annual temperate variation may impact ectotherms’ thermal performance as much or more than predicted changes in average temperature. While suggestive, theoreticians and practitioners alike are still far from understanding the many ways that increasingly variable environmental conditions impact the ability of populations to persist.

Theory points to two ways in which environmental variability affects population fluctuations. The first mechanism is through changes in a population’s long-term average growth rate through nonlinear averaging. Thus, assuming fixed growth under average conditions, increasing environmental variability lowers long-term population growth [11], and the same nonlinear averaging effects apply at the level of demographic rates (see Real and Ellner [12] for a graphical application of this idea). This effect has been demonstrated empirically in the growth rate of experimental populations of green algae (*Tetraselmis tetrahele*) [13], in herbivore feeding rates [14], and in predator feeding rates [15] (although, in some cases, environmental variability can elevate average population growth, especially when the environment has some predictability [16] or populations are spatially distributed [17, 18]).

The second-way environmental variability puts species at greater risk of extinction is by causing short-term population fluctuations. To a first approximation, greater fluctuations in population size enhance extinction risk because populations dip to lower numbers more frequently. At low numbers, populations are at heightened risk of extinction from the increased chance of below-average survival and reproduction among the remaining individuals (i.e., demographic stochasticity; [19, 20]). Taken together, current theory predicts that increases in environmental fluctuations in single-species population models will always drive increases in the long-run population variance [21, 22, 23].

Stochastic population models can predict extinction in both stable [24] and declining populations [25], but do so by assuming relatively simple forms of density-dependence. By simple forms of density-dependence, we mean that which leads to a single stable equilibrium, as in the logistic model with a carrying capacity. Under these models, populations may approach and persist at this equilibrium indefinitely or, below a critical reproductive rate, decline to extinction. This behavior is found in many commonly used models in ecology beyond the logistic model, including the Gompertz, Beverton-Holt, and theta-Ricker models [26, 27]. Ecologists have made progress integrating the effect of environmental variability into our understanding of population and community dynamics that exhibit a single equilibrium [28]. However, these models are ill-suited to describe populations that exhibit more complex dynamics.

In many biological systems, population biologists have recognized the need for models that include more complex forms of density-dependence that generate, for example, multiple equilibria. For example, systems with tipping points between alternative stable states are one case where models of simple density-dependence are insufficient. These systems can exhibit a much richer portfolio of responses to environmental variability than simple density-dependence. For example, in models of alternative stable states, environmental variability can push such a system between these alternative stable states [29], a phenomenon termed stochastic flickering [30, 31]. In addition, stochastic resonance, a sudden increase in system variation, can arise when increases in the extrinsic variance push the system to periodically cycle between alternative stable states when it would otherwise stay in one of the states [32]. This resonance has been hypothesized to play a role in seasonal influenza outbreaks [33] and in the apparent periodic behavior of ice ages [34]. Flickering and resonance predict sudden jumps in the system variance in response to small increases in the environmental variance.

Here, we explore how the variability in the abundance of cycling populations responds to changes in environmental variability. Population cycles are common in nature and serve as a valuable lens for studying dynamical systems with complex feedbacks. We start by analyzing a stochastic, time-dependent Ricker model, which is sufficiently flexible to model both simple density-dependence and cycling populations in a single framework under two cycling mechanisms (following [35]): exogenous cycles that are caused by external factors (e.g., weather [36]) and endogenous cycles that are caused by delayed density feedback loops that can be thought of as self-regulating mechanisms (e.g., stage structure; [37, 38]) or interpopulation mechanisms (e.g., predator effects; [39] or pathogen effects; [40]). Differences between simple density-dependence and density-dependence in cycling populations are visualized in Fig. 1a-c. The most crucial feature is how the qualitative features of density-dependence (illustrated by the contours of the surface) change with density. We use simulations (illustrated in Fig. 1d-f) to qualitatively describe how these models respond to increases in environmental variability and quantify how variation in the environment interacts with the density-dependent dynamics using a novel variance partitioning analysis. We apply this partitioning framework to empirical examples of delayed density dependence in experimental populations of flour beetles (*Tribolium castaneum*) and in predator-prey cycles with a Canadian lynx (*Lynx canadensis*) - snowshoe hare dataset (*Lepus americanus*).

**Figure 1:**
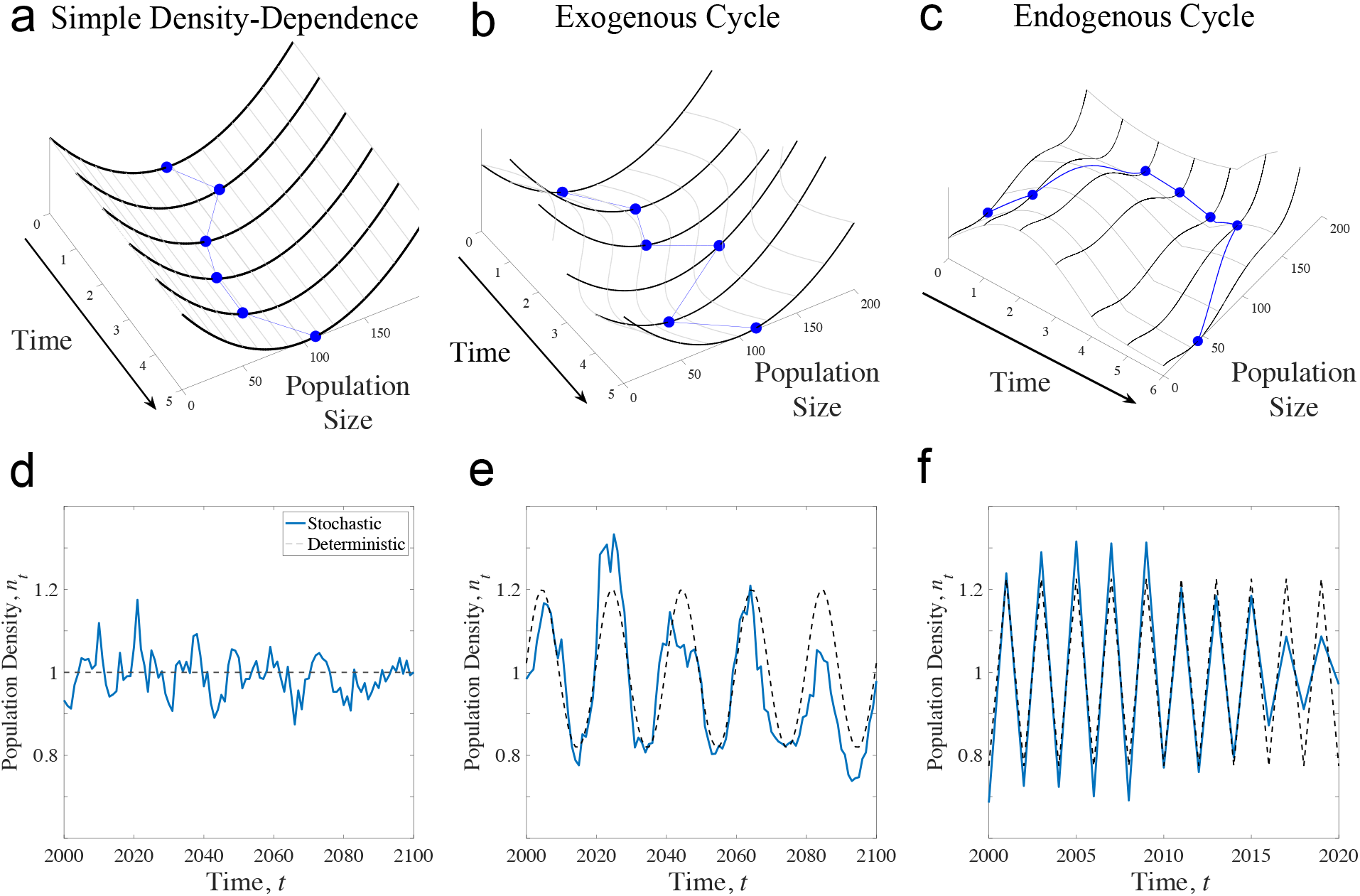
Conceptualization of environmental variability on population dynamics in different models. Under simple models of density-dependence (a), population fluctuations reflect a ball (blue lines) bouncing in a bowl (heavy black lines), where the depth of the bowl (z-axis in all plots) reflects the action of density-dependence. Environmental variation drives random perturbations of the ball away from the bottom of the bowl, which is the carrying capacity in this framework. A conception of exogenous cycles (b) is that the position of the bottom of the bowl moves with the exogenous driver, but the shape of the bowl is unchanging. A conception of endogenous cycles (c) is that the shape and action of density-dependence (black lines) changes over time, meaning that when the population is at low density, the equilibrium moves to higher density, but at high density, the equilibrium moves again to lower density. The topography of the landscape of population change – the bowl itself – is thus changing with the state of the population.

### The impact of variable environments on cycling populations

We start by investigating the properties of the Ricker model, originally developed to qualitatively account for density dependence in the young members of a population [41], for example due to predation. The model has since been derived under additional scenarios [42, 43, 44], and its general applicability to a wide array of empirical conditions has led the Ricker model to be used by many fisheries and wildlife agencies as a management tool. Aiding the adoption of this modeling framework in applied ecology is the ease with which estimation and forecasting can be done, even when observations are contaminated with measurement error [45]. The choice to use the Ricker here, in addition to its wide application in fisheries and wildlife management, is the fact that multiple dynamical regimes can be explored in a single, simple model where environmental variability can be easily incorporated. This allows for direct comparisons of the effect of environmental variability in different dynamical regimes without having to make comparisons between different modeling frameworks.

If *n*_*t*_ is the density of individuals in the population at time *t*, the Ricker model defines the density of individuals in the following time step, *n*_*t*+1_, as 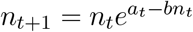. Here, *a*_*t*_ is the (possibly time-dependent) density-independent per-capita growth rate while *b* is a coefficient scaling the effect of density. A typical interpretation is that the species under consideration has discrete generations, and the time *t* is units of generations such that each individual produces 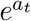 offspring in year *t* and 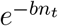 is the fraction that survive.

The Ricker model can exhibit either a single stable equilibrium, exogenous cycles, or endogenous cycles depending on the time-dependent parameter *a*_*t*_. A stable equilibrium occurs when *a*_*t*_ is a constant less than 2 for all *t*. Exogenous cycles occur when *a*_*t*_ is a time-dependent cyclic function (for the simulations below, we chose a sine function), and endogenous cycles occur when *a*_*t*_ is a constant slightly larger than 2, which models overcompensatory density-dependence [e.g., 42, 46]. Examples of the three dynamical regimes are given in Fig. 1d-f. The parameter *b* scales population density without affecting dynamical regimes and so can be re-scaled out of the model; see SI S2.2 for details. The stable-Ricker and similar models have been used to to understand the effects of environmental variability on populations [e.g., 47, 48].

To include environmental variation into the Ricker model we assume that the per-capita growth rate includes random variation over time:

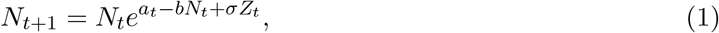

where *Z*_*t*_ is a random variable representing variable environmental conditions in year *t*, and *σ* is a measure of the magnitude of environmental variation over time. Following standard modeling assumptions, *Z*_*t*_ is an i.i.d. standard normal random variable (i.e., *Z*_*t*_ ∼ 𝒩 (0, 1) for all *t*, and *E*[*Z*_*t*_*Z*_*s*_] = 0 for all *t* ≠ *s*). The effect of the environment is given by the term *σZ*_*t*_, which has variance *σ*^2^.

To understand the impact of environmental variation on cycling population dynamics we first consider a small variance approximation [49, 11] in the case of exogenous cycles, then show how this approximation breaks down in the case of endogenous cycles. Small variance approximations simplify the shape of density-dependence around a point (typically an equilibrium) and, as such, describe population dynamics with a linear forms of density-dependence near this point.

#### Exogenous cycles

The approximation for variance in population size in the stochastic Ricker model with an exogenous driver is

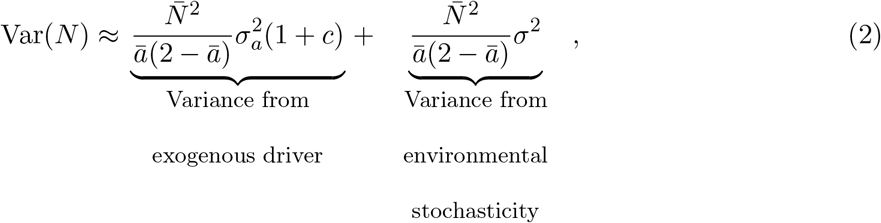

where 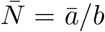 is the average population size (details in S1), *ā* is the average low density growth rate, 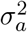 is the (approximate) variance when *a*_*t*_ cycles exogenously, *σ*^2^ is the variance in the stochastic component of the environment, and 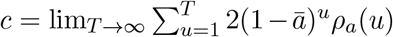 is the effect of autocorrelated environmental variation on population fluctuations (*ρ*_*a*_(*u*) is the autocorrelation function of *a*_*t*_ with time lag *u*; full derivation in supplementary material S3). Equation (2) states that the long-run variance in population size can be partitioned into a sum of two components: variability from a periodic exogenous factor 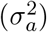 and from environmental stochasticity (*σ*^2^). Approximation (2) predicts a linear increase in population variability with increases in either the periodic exogenous factor or in the random environmental stochasticity. This prediction matches well the observed qualitative linear increase in Var(*N*) with increasing ^2^ for both models of standard density-dependence when 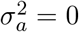(Fig. 2a) and exogenous cycles when when 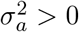 (Fig. 2b). Moreover, this approximation does a good job matching the quantitative relationship between Var(*N*) and 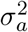 (Fig. S3) and *σ*^2^ (Fig. S4) in the model with exogneous cycles.

**Figure 2:**
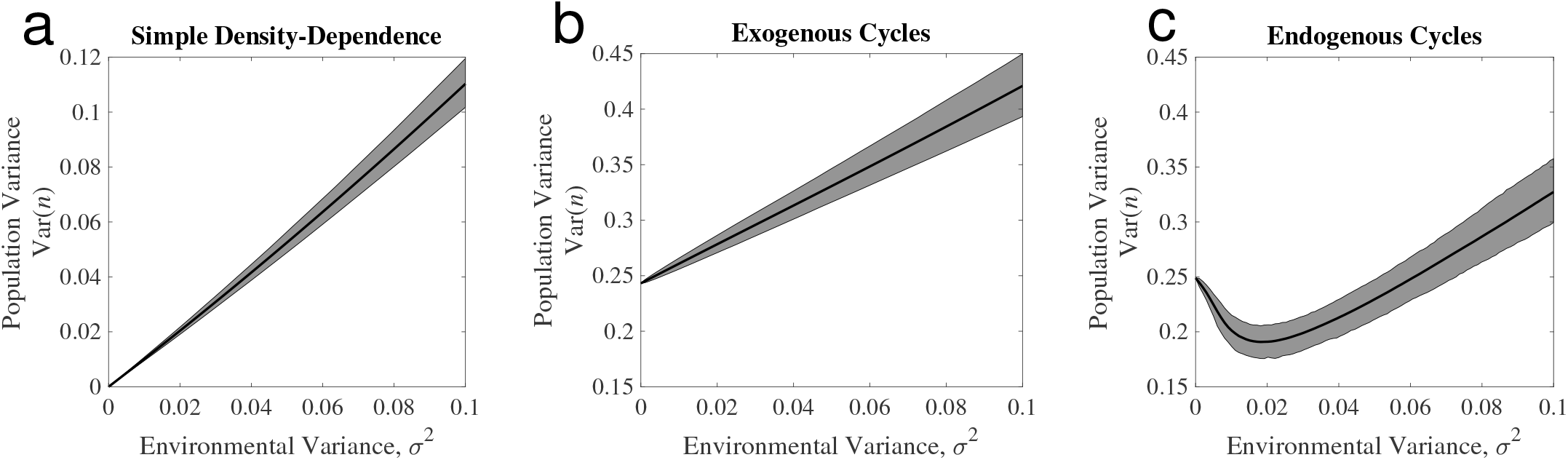
Effect of changing environmental variability for three qualitatively different kinds of dynamics. Each plot shows the average variance of population size over the time series across 1000 replicates (solid black line) as well as the the 95% upper and lower quantiles across replicates. Parameters: In (a), 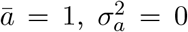, and *b* = 1. In (b), 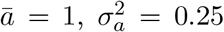, and *b* = 1. In (c), 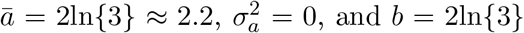, and *b* = 2ln*{*3*}*. The value of *a* in (c) is chosen so that Var(*N*) in the deterministic cycling models are approximately the same, thereby facilitating comparisons of the two different mechanisms of cycles (details in S2).

#### Endogenous cycles

The small variance approximation fails in the endogenous cycle case as eq. (2) predicts negative variance when *a*_*t*_ is a constant greater than 2 (and so *ā* > 2). Simulations of the stochastic endogenously cycling model show that the variance in population size initially declines as the environment becomes more variable (Fig. 2c). The implication here is that, for some levels of environmental variation, increasing environmental variability reduces population fluctuations.

The shape of density-dependence suggests a possible mechanism of this paradoxical decline in population variability with increasing environmental variability. One way to understand density-dependence for the 2-cycle is to consider the second iterate map [50]. If *F* (*n*_*t*_) is the deterministic Ricker map (i.e., *n*_*t*+1_ = *F* (*n*_*t*_)), then the second iterate map is *F*_2_(*n*_*t*_) = *F* (*F* (*n*_*t*_)) which projects population size 2 time-steps in the future. For a 2-cycle, the second iterate map has 3 non-trivial fixed points: 2 stable fixed points corresponding to the two asymptotic cycling values (illustrated with the dashed lines in Fig. 3), and an unstable fixed point exactly between the two (Fig. S1). This unstable fixed point is given by the value *ā/b* (supplementary material S2), and has weak local stability as measured by the local rate of change in the 2-step per-capita growth rate (i.e., the slope of ln*{n*_*t*+2_*/n*_*t*_*}* with respect to *n*_*t*_ at each fixed point). Populations that begin near this weak unstable fixed point can take a long time before exhibiting asymptotic dynamics (Fig. S2), a phenomenon termed a long transient dynamic [51].

**Figure 3:**
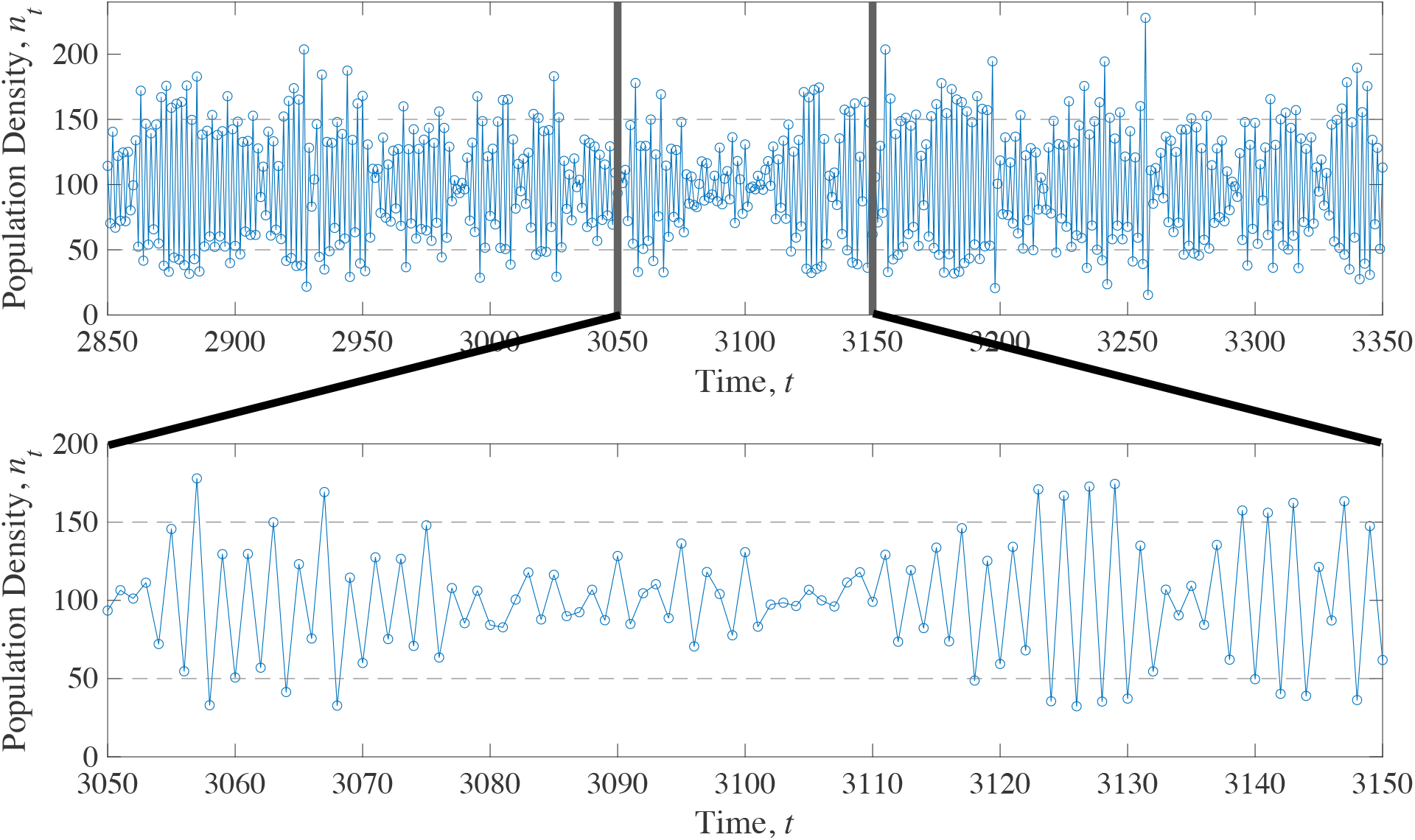
Sample path for the stochastic Ricker model in the case of endogenous cycles showing the origin of stochastic variance induced population fluctuations reduction. Periodically, the population gets “stuck” near the unstable equilibrium separating the two points of the deterministic cycle. In this example, the deterministic model cycles between the two fixed points *n*_*t*_ = 50 and *n*_*t*_ = 150 (dashed lines). The mean population density (here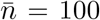) is an unstable fixed point. In this example *a* = 2ln*{*3*}, b* = *a/*100, and *σ*^2^ = 0.02.

The fact that the system can take a long time to escape the unstable fixed point means that the dynamical consequences of stochastic variation depend on both the strength of environmental fluctuations and on the nature of density-dependence. Near stable fixed points, stochasticity has minimal effects due to strong density-dependence. Near the weak unstable fixed point, stochastic effects are relative strong in comparison with the weak effect of density-dependence. To visualize this effect, consider the sample path of the stochastic endogenously cycling model in Fig. 3. Occasionally, stochastic variation pushes the population towards the unstable fixed point (located at 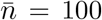in Fig. 3). As a consequence, the population gets temporarily “stuck” and spends extended time in a transient state where fluctuations in population size are mostly stochastic, until the population eventually exits the weak transient phase. Exit of the transient can occur either because of the slow deterministic dynamics back to the cycle or due to large chance perturbations. This dynamical phenomenon keeps the population near the long-run average (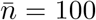in Fig. 3), thereby reducing overall population size variability in comparison to the system without environmental stochasticity. The inset of Fig. 3 shows one particularly acute example of this effect in a representative sample path of the stochastic Ricker model with endogenous cycles.

### Variance decomposition of time series

To quantitatively assess the roles of stochasticity, deterministic dynamics, and their potential interaction, we partitioned the overall variance in population fluctuations (denoted as Var(*N*_*t*_)) as follows:

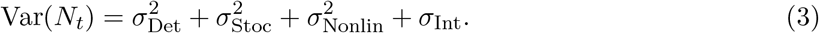

This formula partitions the variance in population size as a sum of four components: (i) the variance in population size in a purely deterministic model, 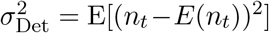, where *n*_*t*_ is the predicted deterministic trajectory in the absence of stochasticity, (ii) the contribution of stochastic variation on top of the deterministic variation, 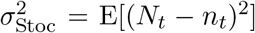, where *N*_*t*_ is the observed stochastic process, the effects of stochasticity on average population density caused by nonlinear averaging (i.e., mean shifts from Jensen’s inequality) 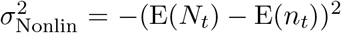, and (iv) and covariation between stochastic and deterministic variation, *σ*_Int_ = 2E(*n*_*t*_ − E(*n*_*t*_))(*N*_*t*_ − *n*_*t*_). The interaction component *σ*_Int_ can be positive or negative, which differs from all other components, which are all strictly non-negative. Its sign measures whether the direction of deterministic fluctuations (*n*_*t*_ − *E*(*n*_*t*_)) are on average in the same direction as stochastic fluctuations (*N*_*t*_ − *n*_*t*_). Thus, *σ*_Int_ *>* 0 when environmental stochasticity amplifies deterministic fluctuations while *σ*_Int_ *<* 0 when environmental stochasticity damps deterministic fluctuations. The derivation of eq. 3 is provided in supplementary material S4.

We applied eq. (3) to simulations from the Ricker model under simple density dependence, exogenous cycles, and endogenous cycles. We found that the Ricker behaves as expected under simple density-dependence and exogenous cycles with *σ*_Int_ = 0. Under endogenous cycling, *σ*_Int_ *<* 0 (Fig. 4), reflecting damping of deterministic fluctuations by environmental stochasticity. This demonstrates that the variance partition accurately measures the novel variance damping behavior shown in the relationship between population variability and environmental variability for cycling populations (Fig. 2c).

**Figure 4:**
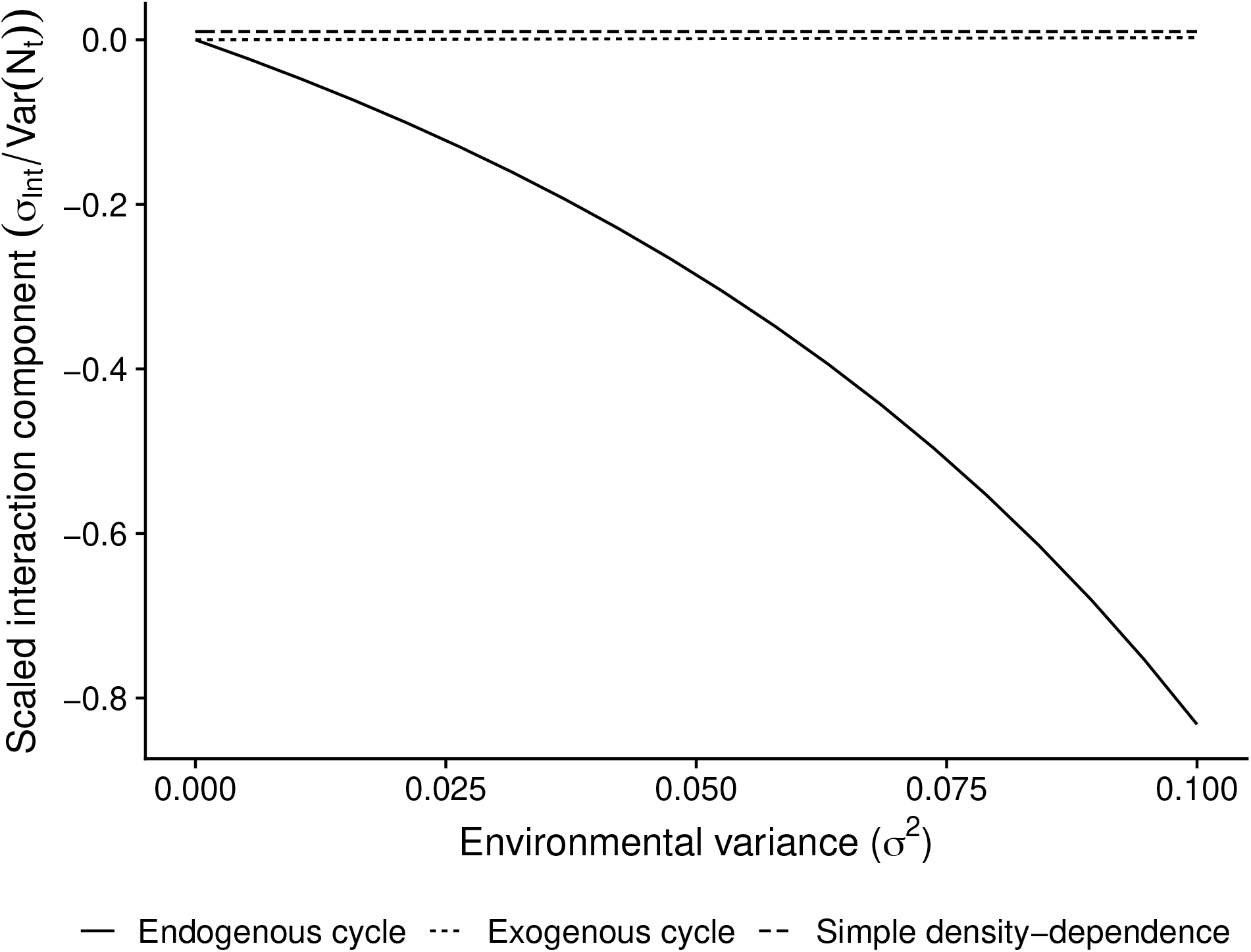
Interaction between environmental variability and deterministic processes as measured by the interaction covariance *σ*_Int_ (eq. [3]) for three different cases of the Ricker model. In the simple density-dependence model, *a*_*t*_ = 1. In the exongenously cycling model, *a*_*t*_ = 1 + sin(*t/π*). In the endogenous cycle mdoel, *a*_*t*_ = 2.2. The interaction term *σ*_Int_ is rescaled by the total population variance Var(*N*_*t*_) to provide a relative measure of the strength of the effect. The lines for simple density-dependence and the exogenous cycle lie exactly on top of each other but are offset slightly in the figure for clarity.

#### Empirical example of lagged density dependence: flour beetle population dynamics

We applied the variance partitioning framework to cyclic populations of flour beetles to test for any evidence of interactions between density-dependence and environmental stochasticity. Experimental data were reported in [52] and are illustrated in Fig. 5. We fit the stage-based larvae-pupae-adult (hereafter LPA) stochastic model described by [53], given as

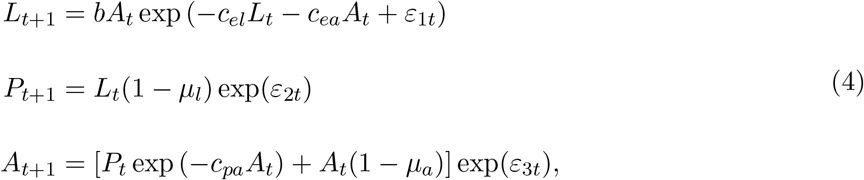

Here, the *ε*_*it*_’s are random values drawn from a multivariate normal distribution where the *ϵ*_*it*_’s are all i.i.d. over time and across life stages. While the LPA model is not the Ricker model, it does have similar qualitative behavior, exhibiting cycles and chaos in some regions of parameter space due to lagged density dependence [52].

**Figure 5:**
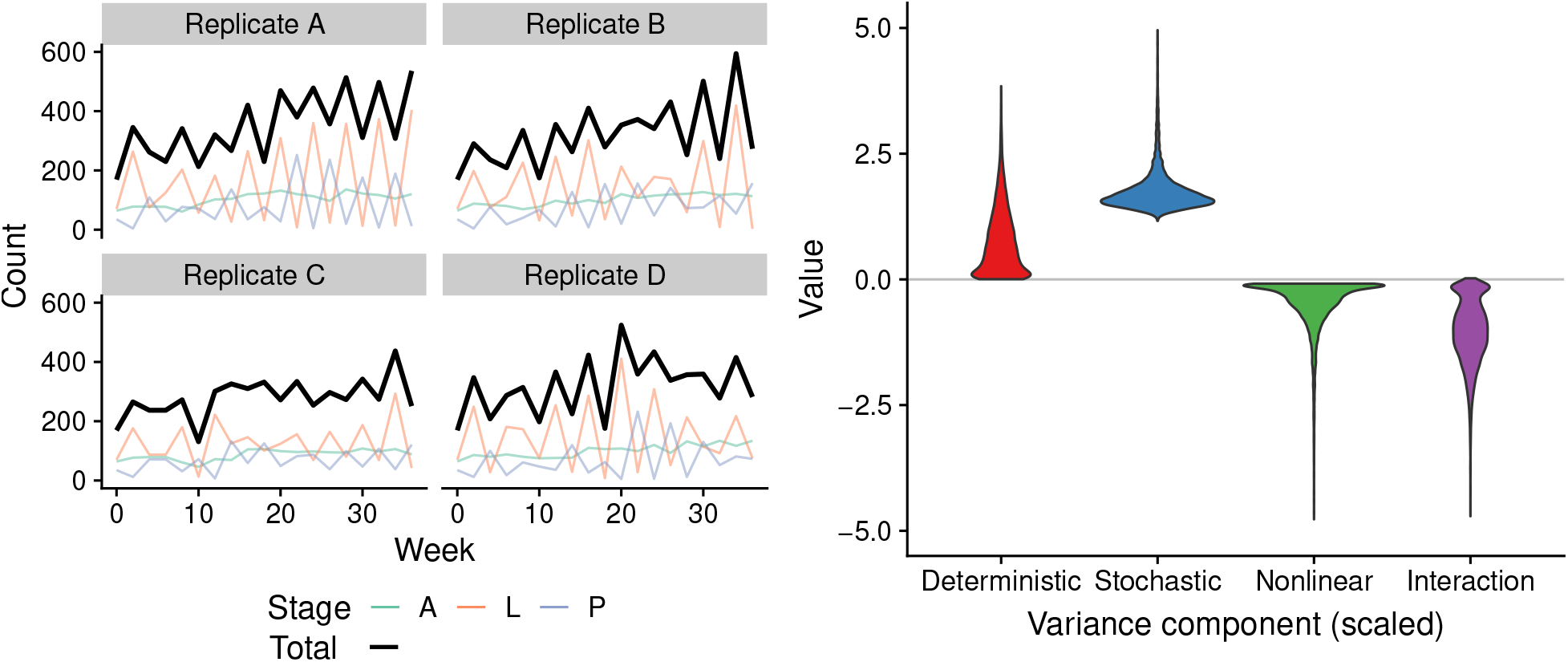
Data for the flour beetle experiment reported in [52] (left panels). Colored lines give weekly counts of each stage, black line gives the total count (*L* + *P* + *A*) used in the variance component analysis. Estimated variance components from eq. (3) (right panel), scaled by the overall population variance.

All parameters were estimated with maximum likelihood using one-step predictions following the methods described in [53]. We used the fitted model to estimate the variance components for the total population abundance, *N*_*t*_ = *L*_*t*_ + *P*_*t*_ + *A*_*t*_, in eq. (3). Further details on estimating the variance components are given in S4.1. Uncertainty in each component was estimated using a parametric bootstrap with 10,000 draws from the sampling distribution. To combine the estimated variance components from each of the four experimental replicates, we used the sum of the variance components of each replicate for each bootstrap draw, which assumes replicates are independent. Using our partitioning scheme, we estimated that variance reduction from interactions between environmental variability and density-dependence reduced population fluctuations by 50%. The sampling distributions of each variance component (reported as mean [95% confidence interval]), normalized by the overall population variance, are shown in Fig. 5. We estimated population fluctuations arising from complex density-dependence alone 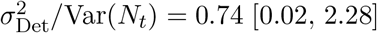, population fluctuations from stochasticity 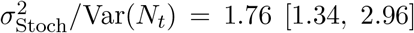, moderate effects of nonlinear Averaging 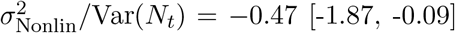, and variance damping sufficient to cancel more than half of the stochastic population fluctuations *σ*_Int_*/*Var(*N*_*t*_) = −1.02 [-2.4, -0.1]. The interaction estimate indicates that the covariance between stochastic and deterministic dynamics plays a sizeable role in damping the overall variance, and that, without this damping effect, the overall population variance would be about double what is observed.

#### Empirical example of predator-prey dynamics: lynx-hare interactions

While intraspecific interactions are one mechanism that generates endogenous cycles, trophic interactions, such as predation, can also create population cycles. Here, we used ninety-one years of data on Lynx and snowshoe hare (*Lepus americanus*) pelts from the Hudson Bay Company (data from [54]) to model predator-prey dynamics. We then applied the variance partitioning framework to the fitted model of the Canadian Lynx to estimate the interaction component in eq. (3). We fit a discrete-time version of the Lotka-Volterra predator-prey model where Lynx (*L*) and hare (*H*) dynamics follow

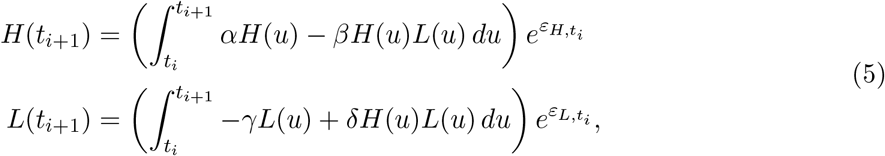

where the 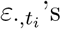 are random values drawn from normal distributions that are i.i.d over time and species. Parameters from eq. (5) were estimated using maximum likelihood in the template model builder R package [55]. We assumed observations of hare and Lynx abundance (Fig. 6A) were lognormally distributed. We also assumed that the first observations in the time series were drawn from a lognormal distribution with unknown mean and the same variance as subsequent observations. Using the fitted model, we estimated the variance components and their uncertainty in the same manner as above. Further details on the estimation of the variance components and their uncertainty are given in S4.1.

**Figure 6:**
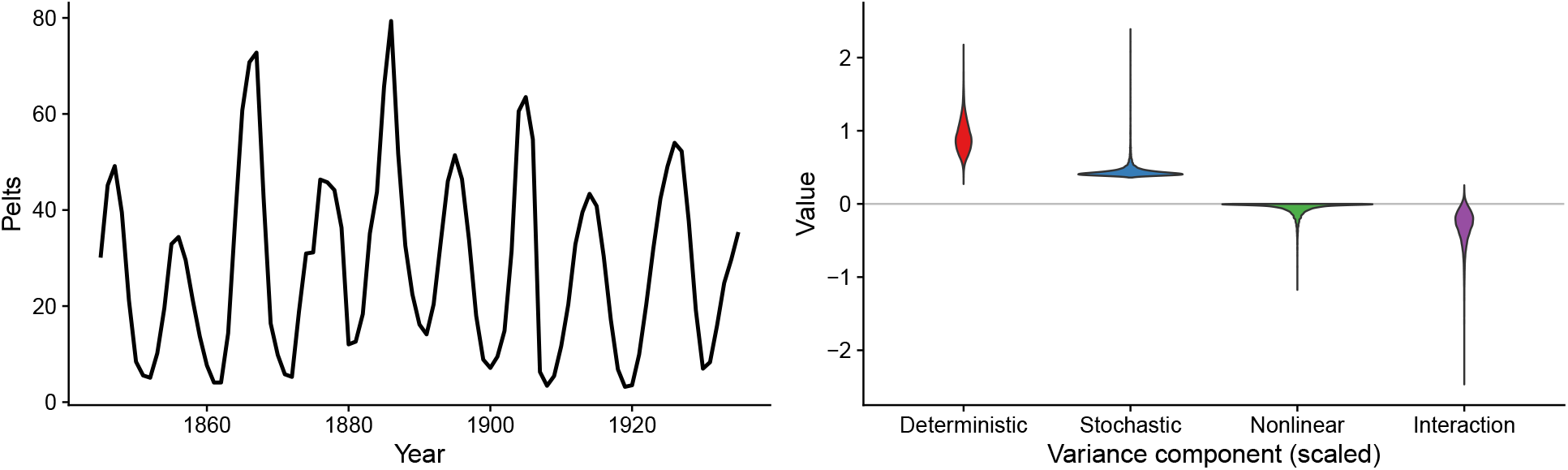
Lynx population data (left panel). This times series was modeled using eq. (5). Estimated variance components from eq. (3) (rescaled by the overall population variance) are illustrated in the right panel.

As with the flour beetle case studied above, we find again that the interaction between density-dependence and environmental variability has effects on population fluctuations of similar orders of magnitude to either operating alone. The sampling distributions of each variance component, normalized by the overall population variance, are shown in Fig. 6. We estimated a dominant contribution of deterministic dynamics to overall population fluctuations 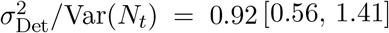, smaller effects of environmental stochasticity 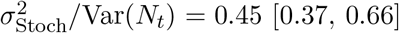, weak nonlinear averaging 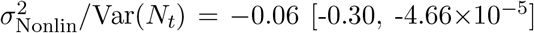, and sizeable variance damping *σ*_Int_*/*Var(*N*_*t*_) = −0.31 [-0.92, 0.03]. Here, we estimate that population fluctuations would be ≈ 31% more variable without the interaction between complex density-dependence and environmental fluctuations.

## Discussion

As global change continues, one expectation is that conditions will become more variable over time. With increasing environmental variability, a common worry is that many populations will experience greater abundance fluctuations and a corresponding increase in the probability of extinction. Such a perspective derives from models assuming populations fluctuate around a steady state and that increasing variability in the environment will drive increases in population fluctuations [56, 57, 58, 47]. Here, we considered the response of non-equilibrium dynamics, namely cycles, to increasing environmental variability. We found that increasing environmental variability in an exogenous cycle (i.e., caused by a dynamically unlinked cycling factor) leads to proportional increases in population fluctuations, as expected from prior models assuming simple density dependence (i.e., a point equilibrium). However, endogenous cycles driven by lagged density-dependence respond to environmental variability qualitatively differently; increasing environmental variability can diminish overall population fluctuations. The mechanism diminishing population fluctuations reflects an interaction between environmental variation and density-dependence that relies on the presence of long-transients in the deterministic system. A long transient is a dynamic that is not represented in the asymptotic behavior of a dynamical system, but lasts long enough to be relevant on the timescales of typical ecological observations [51].

In many domains of ecology and evolution, researchers have recognized the potential for stochastic factors to interact in non-intuitive ways with deterministic processes. Coulson et al. [59] cataloged a number of notable examples of these interactions in ecology and epidemiology and subsequent work has noted additional ways environmental variability can interact in complex ways with deterministic dynamics. Nearly all these studies have emphasized how environmental variability can amplify deterministic fluctuations, or generate new fluctuations (i.e., a positive interaction in eq. [3]). For example, environmental variability has been shown to push a system currently in one stable state to another [29] or to “stochastically flicker” between alternative stable states when it would otherwise remain in one of the stable states [30]. In addition to stochastic flickering, stochastic resonance [32] and stochastic amplification [60] both describe interactions between stochastic variation and deterministic dynamics that increase system variability. Our study shows that these examples fall into a larger class of interactions that also include the paradoxical phenomenon whereby increases in stochastic variation reduce overall system variability.

The variance damping explored here appears to be most similar to a phenomenon noted in the literature as inverse stochastic resonance. As first described in models of neural circuits, the oscillatory firing of neurons collapses to a fixed point with increases in stochastic variation [61]. Touboul et al. [31] also described inverse resonance arising in an ecosystem model of savanna-forest dynamics. Another potential example of this phenomenon may be related to work on whooping cough, where Rohani et al. [62] found that increasing levels of stochasticity in disease models slowed down annual epidemic cycles. More work needs to be done to determine if the mechanism driving all of these phenomenon is equivalent to the damping process we describe here. In any case, all would be undoubtedly classified as negative interactions in our variance partitioning scheme, and together suggest an interesting avenue for future research and synthesis.

The partitioning scheme we develop has broad applications to any ecological system in which a deterministic trajectory can be described and researchers want to understand the relative contributions of stochastic and deterministic factors to population fluctuations. Recent work by Mutshinda et al. [63] determined the contribution of density-dependent dynamics to the long-run population variance by comparing the predicted long-run variance to a density-independent model under the same stochastic forces. The partitioning scheme we present here (eq. (3)) addresses a similar problem, but differs in that it allows for, and quantifies, interactions between density-dependence and stochastic variation. In addition to the interaction, the partitioning can be interpreted in terms of three well-known terms: (i) the deterministic dynamics that can include population cycles, (ii) the stochastic dynamics that account for the complex sources of variation that appear unstructured, and (iii) the differences between the observed average population density and the average predicted by a deterministic model arising from Jensen’s inequality.

Biologists have long relied on variance partitioning schemes to identify components of variation that have conceptual value, even if their underlying mechanistic causes are obscured. For example, analysis of variance, one of the most commonly applied statistical tools in biology, is used to distinguish differences among and between groups by comparing the average within group variation to the between group variation. Similarly, quantitative genetics has successfully partitioned the additive heritable phenotypic variation into genetic and environmental components without knowing the specific loci responsible for this variation [64]. The study of competition and species coexistence now routinely uses a partitioning scheme that envisions counterfactual scenarios (e.g., a system in which no stabilizing mechanisms are present) to measure the impact of species differences in maintaining or eroding species diversity [65, 66]. Variance partitioning schemes for population models can play a similar role, helping us to categorize and quantify the qualitative components of a dynamical system of very different systems and improve our understanding of the mechanisms that drive fluctuations in population size over time. As a descriptor, this partition is useful for identifying, measuring, and categorizing drivers of population variation in nature and in models. We expect that a wide-scale and data-driven analysis of variance components applied to population time-series should reveal the prevalence and evidence for variance damping in nature.

Understanding the many impacts of global change on species demands an approach that embraces variability and uncertainty. Simple population models with environmental stochasticity are mathematical microscopes (sensu Cohen [67]) of natural systems through which we infer how population processes work under such uncertainty and variability. Recent analyses suggest that the prevalence of nonlinear dynamics may be much more common than previously appreciated [68, 69], so understanding the stochastic behavior of the models with nonlinear dynamics, especially relatively simple ones such as the Ricker model, is an important step toward making sound forecasts about the future of these populations. Our results suggest that our baseline understanding of environmental variability in complex ecological systems may need to be revised. Even elementary modifications to simple models may reveal paradoxical, yet plausible, phenomena such as the variance damping explored here. Our investigation into variance damping in flour beetle and Lynx-Hare dynamics shows that more complex, biologically grounded models may have similar paradoxical behavior. Given the inferential and predictive roles that stochastic population models play in applied population and conservation biology [70, 71], understanding the basic dynamical properties of commonly used models remains imperative. As such, damping interactions between environmental stochasticity and deterministic dynamics may play a previously unrecognized role in the dynamics of ecological systems.

## Acknowledgements

Many thanks to Bob Holt and the Holt lab group for comments on previous versions of this manuscript. Funding was provided by the National Science Foundation grant NSF-DMS-2052372 to JMF and JMP.

## Code Statement

Code to reproduce the work found in this manuscript can be found at https://github.com/kortessis/Nonlinear-Dynamics-and-Environmental-Variability.

## Competing Interests Statement

The authors declare no competing interests related to this work.

## Notes

https://github.com/kortessis/Nonlinear-Dynamics-and-Environmental-Variability

### Competing Interest Statement

The authors have declared no competing interest.

### Summary of Updates

Potential functions removed and replaced with empirical demonstration of variance dampening using flour beetle and Lynx-Hare time series data.

