## Supplementary materials for "Increasing environmental fluctuations can dampen variability of endogenously cycling populations"

### Supplementary information for ”Increasing environmental fluctuations can dampen variability of endogenously cycling populations”

[1], Nicholas Kortessis*[1,2], Jake M. Ferguson[3], and José Miguel Ponciano[2]

[1]Department of Biology, Wake Forest University
[2]Department of Biology, University of Florida
[3]Department of Biology, University of Hawaii at Manoa

### S1 Mean population size of the Ricker model

Here, we determine the mean population size of the Ricker model, which is needed to calculate the variance. Consider the stochastic model (equation [1] of the main text) with $\sigma$ as the measure of environmental variability. The deterministic model can be recovered by simply taking $\sigma = 0$ in the following expressions.

Consider $\{N_t\}$, the sequence of time-dependent random variables of population size in the Ricker model. The value of the $t$-th random variable is given by the one-step recursion

$$N_t = N_{t-1} e^{a_{t-1} - bN_{t-1} + \sigma Z_{t-1}} \tag{S1}$$

where random variables are denoted with uppercase letters and known values at a specific time (e.g., $a_t$) are denoted with lowercase letters. Taking logs of both sides gives

$$X_t = X_{t-1} + a_{t-1} - bN_{t-1} + \sigma Z_{t-1}, \tag{S2}$$

where $X_t = \ln N_t$ is population density on the log scale. We can rewrite this expression as a function of population sizes two time steps in the past, i.e.,

$$X_t = X_{t-2} + \sum_{i=t-2}^{t-1} a_i - b \sum_{i=t-2}^{t-1} N_i + \sigma \sum_{i=t-2}^{t-1} Z_i \tag{S3}$$

Extending this process back to the initial time step, $t = 0$, it follows that the population size at time $t$ (on the log scale) is a sum of four terms:

$$X_t = X_0 + \sum_{i=0}^{t-1} a_i - b \sum_{i=0}^{t-1} N_i + \sigma \sum_{i=0}^{t-1} Z_i, \tag{S4}$$

where the first term is the initial state of the population, the second term is the cumulative effect of density-independent growth, the third term is the effect of density-dependence, and the final term is the effect of environmental stochasticity.

Dividing through by $t$ and taking the limit as $t \to \infty$ shows that the effect of initial conditions decay from the system such that

$$0 = \lim_{t\to\infty} \frac{1}{t}\sum_{i=0}^{t-1} a_i - \lim_{t\to\infty} \frac{1}{t} b \sum_{i=0}^{t-1} N_i + \sigma \lim_{t\to\infty} \frac{1}{t}\sum_{i=0}^{t-1} Z_i \tag{S5}$$

because $\lim_{t\to\infty} X_i/t = 0$ for any bounded population $X_i$, i.e., $|X_i| < \infty$.

Note in equation (S5) that each limit is a long-term average, whether the average is of a sequence of deterministically varying processes (such as $\{a_t\}$) or a sequence of random variables (such as $\{Z_t\}$). To deal with the random environmental variables, recognize that $\sum_{i=0}^{t-1} Z_i$ is a sum of $t$ independent standard normals. A standard result in statistics is that such a sum is itself normally distributed with mean 0 and variance $1/t$. Hence, in the limit of large $t$, the sum converges uniformly on the value 0, i.e., $\lim_{t\to 0} t^{-1}\sum_{i=0}^{t-1} Z_i \to 0$.

Substituting in $\lim_{t\to 0} t^{-1}\sum_{i=0}^{t-1} Z_i \to 0$ and solving for the long-term mean of $N$ yields

$$\bar{N} = \lim_{t\to\infty} \frac{1}{t}\sum_{i=0}^{t-1} N_i = \frac{1}{b}\left[\lim_{t\to\infty} \frac{1}{t}\sum_{i=0}^{t-1} a_i\right] = \frac{\bar{a}}{b}, \tag{S6}$$

where an overbar indicates the long time average of a time-varying quantity. In particular, $\bar{a}$, the long-term average maximum growth rate, plays a central role in the approximations developed in section S3 below.

Expression (S6) holds provided that $b > 0$, i.e., the model is density-dependent, and that the long-term average of $\{a_t\}$ is finite.

### S2 Variance of the deterministic Ricker model

As a baseline against which to compare the effects of environmental variability, we need to determine the variance of the deterministic Ricker model in the different cases. Naturally, the long-term variance of the case with a stable equilibrium is zero because the population does not change at the equilibrium in the deterministic model. We focus on the variance of the exogenous and endogenous cycles.

#### S2.1 Variance of the 2 cycle

To find the variance of the Ricker model exhibiting period-2 cycles, recognize that the two points in the cycle are equilibrium points of the 2-step Ricker map. The 2-step Ricker map can be found by iterating the one-step recursion twice such that population size at time $t + 2$ is given as an explicit function of time $t$, i.e.,

$$n_{t+2} = n_t \exp\left\{2a - bn_t\left(1 + e^{a-bn_t}\right)\right\}. \tag{S7}$$

A representative example of the 2-step map is given in Fig. S1. Any equilibrium of the two-step map satisfies $n_{t+2} = n_t$ and therefore falls along the intersection of the solid and dotted lines in Fig. S1. It is then the case that any (non-trivial) equilibrium point, denoted by a hat, satisfies

$$2a - b\hat{n}\left(1 + e^{a-b\hat{n}}\right) = 0. \tag{S8}$$

One can see immediately that $b\hat{n} = a$ (i.e., $\hat{n} = a/b$) is one solution to (S8). For $0 < a \leq 2$, this is the only solution and is the carrying capacity. At $a = 2$, there is a super-critical bifurcation such that $\hat{n} = a/b$ switches from being globally stable to locally unstable, with the coincident emergence of two other solutions which are each locally stable. The two emergent solutions appear as alternative stable equilibria in the two-step map, but are simply the points that the model oscillates between in the one-step map. Call these solutions $\hat{n}_A$ and $\hat{n}_B$.

Given that there are only two points in the asymptotic set of period-2 cycles and each point is equally frequent over long times, the long-term variance is

$$\lim_{t\to\infty} \mathrm{Var}(n) = \frac{1}{2}\left[(\hat{n}_A - \bar{n})^2 + (\hat{n}_B - \bar{n})^2\right]. \tag{S9}$$

Given that there are only two points in the cycle, it must be the case that the two points are equally distant from the average because an average of two values is simply the midpoint between them. We represent this distance between the mean and a point in the 2-point cycle in relative units of average population size, i.e., define

$$\Delta = \frac{|\hat{n}_j - \bar{n}|}{\bar{n}} \tag{S10}$$

for $j = A, B$. This definition describes the amount of relative variation in the population but removes any effects of population size. To see why, note that if one were to use $\Delta$ in the expression for the variance of the period-2 cycle, the variance has the following remarkably simple form:

$$\begin{aligned}\lim_{t\to\infty} \mathrm{Var}(n) &= \frac{1}{2}\left(\Delta^2\bar{n}^2 + \Delta^2\bar{n}^2\right) \\ &= \Delta^2\bar{n}^2.\end{aligned} \tag{S11}$$

As such, it is the case that $\Delta$ is the asymptotic coefficient of variation,

$$\lim_{t\to\infty} \mathrm{CV}(n) = \frac{\sqrt{\mathrm{Var}(n)}}{\bar{n}} = \Delta, \tag{S12}$$

which measures of variation in population size when standardized by the mean. Hence, $\Delta$ is a non-dimensional quantity representing the amount of variability in the 2-cycle.

While it is analytically difficult to find the value of $\Delta$ for specified values of $a$ and $b$, it is feasible to first choose a value of $\Delta$, and then find the corresponding values of $a$ and $b$. To do so, we rewrite equation (S8) explicitly in terms of $\Delta$, which can be done by substituting $\hat{n}_A = (a/b)(1+\Delta)$ for $n_t$, and simplifying, which yields

$$(1-\Delta) - (1+\Delta)e^{-a\Delta} = 0. \tag{S13}$$

Solving for $a$ gives

$$a = \frac{1}{\Delta}\ln\left\{\frac{1+\Delta}{1-\Delta}\right\}. \tag{S14}$$

Equation (S14) is valid for $0 < \Delta \lesssim 0.722$, which corresponds to $2 < a \approx 2.526$, the region of parameter space corresponding to the 2-cycle.

One may do the same exercise with the definition of $\hat{n}_B$, the smaller value of the 2-cycle and arrive at an expression for $a$ that is identical to expression (S14). For the purposes of simplification in the main text, we have chosen $\Delta = 0.5$, meaning that the population fluctuates between $1.5\bar{n}$ and $0.5\bar{n}$ in the 2-pt cycle. Using $\Delta = 0.5$ in equation (S8) above means that $a = 2 \cdot \ln\{3\}$ and that the variance for the 2-pt cycle is $0.25\bar{n}^2$.

We can also approximate the value of $\Delta$ through a Taylor series expansion of equation (S14). Since this can be rewritten as $\Delta = \frac{2}{a}\sum_{\text{i odd}} \Delta^i/i$, a third order approximation in $\Delta$ gives $\Delta \approx \frac{2}{a}\left(\Delta + \Delta^3\right)$. Thus, the value of $\Delta$ is dependent on $a$ and is given by $\Delta \approx \frac{3a}{4}\sqrt{\frac{8}{3a}\left(1-\frac{2}{a}\right)}$.

#### S2.2 Variance of the exogenous cycle

Calculating the variance of the exogenous cycle is more complicated than the endogenous 2-pt cycle. Unless the exogenous variation is quite simple, it can be prohibitively difficult to get an exact expression for the variance. Rather than trying to get an exact expression for the variance, we resort to an approximation of the deviations of population size from its long-term average, $\bar{n}$, which is sufficient for approximating the variance in population sizes.

The approximation works by writing population size as a linear function of $a_t$ and $n_t$ assuming that neither variable deviates too much from their respective mean values, $\bar{a}$ and $\bar{n}$. One issue that immediately arises is to determine what constitutes a large deviation, since $a_t$ and $n_t$ need not fluctuate on similar scales of magnitude. The scale of variation in $a_t$ is a parameter that we call $\sigma_a$ and can be independently controlled. We assume that $a_t$ has the general form of $a_t = \bar{a} + \sigma_a X_t$ where $X_t$ has mean zero over time and variance over time of 1. For example, a cyclic environmental process, we could use a discretized sine wave function of the form

$$a_t = \bar{a} + \sigma_a\sqrt{2}\sin\left(2\pi\frac{t-\tau}{\Omega}\right) \tag{S15}$$

where $\Omega$ is the period of $a_t$ and $\tau$ depends on the value of $a_0$. For sufficiently large $P$, we may define $X_t = \sqrt{2}\sin(2\pi(t-\tau)/P)$ and note that the long-term average of $X_t$ is zero and the variance is 1. As such, $\bar{a}$ is the mean and $\sigma_a^2$ is the variance of $a_t$.

While variation in $a_t$ can be controlled by a parameter, $n_t$ is a variable that fluctuates according to the variation in $a_t$. We need to standardize $n_t$ to similar order as the variation in $a_t$ to make use of the approximation. One way to standardize population size variation is to rewrite the Ricker map in terms of $n_t' = n_t/\bar{n}$, population density relative to the mean value, $\bar{n} = \bar{a}/b$. Note that the mean value of relative population density is $T^{-1}\sum_{t=1}^{T} n_t' \to (T^{-1}\sum_{t=1}^{T} n_t)/\bar{n} = 1$ for large $T$.

To do the approximation, first label the Ricker map at any time step as

$$n_{t+1}' = F(a_t, n_t') \tag{S16}$$

where $F(a_t, n'_t) = n'_t \exp\{a_t(1 - n'_t)\}$. To first-order approximation, relative population density in time $t + 1$ is

$$n'_{t+1} = F(\bar{a}, 1) + F_{a_t}(\bar{a}, 1)(a_t - \bar{a}) + F_{n'_t}(\bar{a}, 1)(n'_t - 1) + O(\sigma_a^2) \tag{S17}$$

where $F_x(\bar{a}, 1) = \partial F/\partial x|_{a_t=\bar{a}, n'_t=1}$ is the partial derivative of $F$ w.r.t. $x$ evaluated at the equilibrium point $(a_t = \bar{a}, n'_t = 1)$, and $O(\sigma_a)$ is a measure of the order of variability in $a_t$. This "big-o" notation simply indicates that all terms of order $\sigma_a^2$ or higher in the Taylor series expansion can be collected in a single term. Formally, $y = O(x)$ (read "$y$ is big-o of $x$") means that $|y| \leq Cx$ as $x \to 0$ for any constant $C > 0$. Formally, we stipulate that $a_t - \bar{a} = O(\sigma_a)$ for all $t$. This is easily controllable in the sine wave function we consider for exogenous cycles in the main text.

To proceed, we need the coefficients of the Taylor series. Some algebra shows that $F(\bar{a}, 1) = 1$ and some calculus reveals that

$$\begin{aligned} F_{a_t}(\bar{a}, 1) &= 1 \\ F_{n'_t}(\bar{a}, 1) &= 1 - \bar{a}. \end{aligned} \tag{S18}$$

The above Taylor expansion, once explicitly writing the coefficients, rearranging, and taking squares yields

$$(n'_{t+1} - 1)^2 = (a_t - \bar{a})^2 + (1 - \bar{a})^2 (n'_t - 1)^2 + 2(1 - \bar{a})(a_t - \bar{a})(n'_t - 1) + o(\sigma_a^2). \tag{S19}$$

where $o(\sigma_a^2)$ ("little-o" notation) indicates that all terms of higher order than $\sigma_a^2$ are negligible for sufficiently small $\sigma_a$. Formally, $y = o(x)$ means that $y/x \to 0$ as $x \to 0$. Hence, we assume that the approximation applies when, as $\sigma_a \to 0$, all other higher-order terms approach zero faster than $\sigma_a$.

This is the squared deviation in population size from its mean in a single time-step. What we need to get a variance is to take the average of squared deviations for infinite time, i.e.,

$$\lim_{T \to \infty} \frac{1}{T} \sum_{t=1}^{T} (n'_t - 1)^2. \tag{S20}$$

Using (S19) in (S20) yields

$$\begin{aligned} \lim_{T \to \infty} \frac{1}{T} \sum_{t=1}^{T} (n'_{t+1} - 1)^2 = \lim_{T \to \infty} \frac{1}{T} \sum_{t=1}^{T} (a_t - \bar{a})^2 + (1 - \bar{a})^2 \lim_{T \to \infty} \frac{1}{T} \sum_{t=1}^{T} (n'_t - 1)^2 \\ + 2(1 - \bar{a}) \lim_{T \to \infty} \frac{1}{T} \sum_{t=1}^{T} (a_t - \bar{a})(n'_t - 1) + o(\sigma_a^2). \end{aligned} \tag{S21}$$

An issue now is to determine the cross-product term, $(a_t - \bar{a})(n'_t - 1)$, which we cannot get an exact expression for because it requires an exact solution for $n'_t$. This term may not be negligible, in fact, because we expect that the population tracks (with some possible time lag) the moving equilibrium determined by $a_t$, and so it would be ill-advised to assume that there is no statistical relationship between $a_t$ and $n'_t$.

To get a handle on this value, we need the linear approximation for $n'_t$, which from (S17) above is

$$n'_t = a_{t-1} - \bar{a} + (1 - \bar{a})(n'_{t-1} - 1) + o(\sigma_a). \tag{S22}$$

Of course, this is itself a function of $n'_{t-1}$. We apply the same linear approximation to the entire history of $n'$, which yields the following linear approximation of $n'_t$:

$$n'_t \approx \underbrace{(1 - \bar{a})^t (n'_0 - 1)}_{\text{Effect of initial state}} + \underbrace{\sum_{u=1}^{t} (1 - \bar{a})^{u-1} (a_{t-u} - \bar{a} - 1)}_{\text{Effect of environmental history}} . \tag{S23}$$

Now, substituting this approximation for $n'_t$ into the cross-product term yields

$$(a_t - \bar{a})(n'_t - 1) \approx (a_t - \bar{a})(1 - \bar{a})^t (n'_0 - 1) + (a_t - \bar{a}) \sum_{u=1}^{t} (1 - \bar{a})^{u-1} (a_{t-u} - \bar{a} - 1). \tag{S24}$$

In the approximation, we need the long-term average of this cross-product, which is given by taking the average in the the limit of large $T$. In that limit, the RHS of equation S24 simplifies to

$$\lim_{T \to \infty} \frac{1}{T} \sum_{t=1}^{T} (a_t - \bar{a})(n'_t - 1) \approx \lim_{T \to \infty} \frac{1}{T} \sum_{t=1}^{T} \sum_{u=1}^{t} (1 - \bar{a})^{u-1} (a_t - \bar{a})(a_{t-u} - \bar{a}), \tag{S25}$$

provided that $0 \le \bar{a} \le 2$ because $\lim_{t\to\infty}(1-\bar{a})^t/t \to 0$ in this range, but is otherwise unbounded.

The RHS of equation S25 can be interpreted as follows. For any value of $T$, the expression is the covariance between $a$ at time $t$ and and some time lag $u$ for all possible values of $t = 1, 2, \ldots, T$ and all time lags, $u = 1, 2, \ldots, t$. Hence, it is a sum of autocovariances across the entire history of the sequence of $a$. The sum is weighted by coefficients, $(1-\bar{a})^{u-1}$ that represent a geometric series and so give greater weight to shorter time lags than longer time lags.

One way to intuit the biological importance of this term is that it summarizes the effect of the history of $a$ on the current population size. When $\bar{a} \approx 1$, the geometric series diminishes quickly and only a short history is needed to describe the current population size. When $\bar{a}$ deviates substantially from 1—but is still within (0,2)—then longer histories of the process of $a$ are required to explain the current population size.

Placing approximation (S25) into (S21) yields the following expression for the limiting squared deviations in relative population size:

$$\begin{aligned}\lim_{T\to\infty}\frac{1}{T}\sum_{t=1}^{T}(n'_{t+1}-1)^2 &= \lim_{T\to\infty}\frac{1}{T}\sum_{t=1}^{T}(a_t-\bar{a})^2 + (1-\bar{a})^2\lim_{T\to\infty}\frac{1}{T}\sum_{t=1}^{T}(n'_t-1)^2 \\ &\quad + 2(1-\bar{a})\lim_{T\to\infty}\frac{1}{T}\sum_{t=1}^{T}\sum_{u=1}^{t}(1-\bar{a})^{u-1}(a_t-\bar{a})(a_{t-u}-\bar{a}) + o(\sigma_a^2).\end{aligned} \tag{S26}$$

In infinite time, the averages in (S26) converge to variances, in which case (S26) rewrites as

$$\mathrm{Var}(n') = \mathrm{Var}(a) + (1-\bar{a})^2\mathrm{Var}(n') + 2(1-\bar{a})\lim_{T\to\infty}\frac{1}{T}\sum_{t=1}^{T}\sum_{u=1}^{t}(1-\bar{a})^{u-1}(a_t-\bar{a})(a_{t-u}-\bar{a}) + o(\sigma_a^2). \tag{S27}$$

Furthermore, in infinite time, the last term of (S27) is dominated by the long-time sums, which are autocovariances because the cross-product average, $T^{-1}\sum_{t=1}^{T}(a_t-\bar{a})(a_{t-u}-\bar{a})$, satisfies the definition of an autocovariance with time lag $u$ for sufficiently large $T$. Autocovariances are just non-standardized autocorrelation functions. As such, we can rewrite this term in infinite time as a sum of the autocorrelation function of $a$, $\rho_a(u)$, which is defined as

$$\rho_a(u) = \frac{\mathrm{Cov}(a_t, a_{t-u})}{\mathrm{Var}(a_t)} = \frac{\frac{1}{T}\sum_{t=1}^{T}(a_t-\bar{a})(a_{t-u}-\bar{a})}{\frac{1}{T}\sum_{t=1}^{T}(a_t-\bar{a})^2}. \tag{S28}$$

This definition implies that the autocovariance is $\mathrm{Cov}(a_t, a_{t-u}) = \mathrm{Var}(a_t)\rho_a(u)$. As such, we rewrite approximation (S27) as

$$\mathrm{Var}(n') = \mathrm{Var}(a) + (1-\bar{a})^2\mathrm{Var}(n') + 2(1-\bar{a})\mathrm{Var}(a)\sum_{u=1}^{T}(1-\bar{a})^{u-1}\rho_a(u) + o(\sigma_a^2). \tag{S29}$$

Solving (S29) for $\mathrm{Var}(n')$ yields our final result:

$$\mathrm{Var}(n') = \frac{\mathrm{Var}(a)}{1-(1-\bar{a})^2}\left[1 + \underbrace{\lim_{T\to\infty}\sum_{u=1}^{T}2(1-\bar{a})^u\rho_a(u)}_{\substack{\text{Effect of autocorrelation}\\ \text{structure of exogenous driver}}}\right] + o(\sigma_a^2). \tag{S30}$$

Approximation (S30) says that the variance in population density scales linearly with the variance in the exogenous driver, $a_t$, in units of growth rate. In the main text, we simplify this expression by defining $c = \lim_{T\to\infty}\sum_{u=1}^{T}2(1-\bar{a})^u\rho_a(u)$ to represent the effect of autocorrelations in the exogenous driver on population variability. Given that it is an average of autocorrelations, it is bounded between -1 and 1. Figure S3 shows the approximation in comparison to Monte Carlo simulations of the exogenous Ricker model for different values of $\bar{a}$ and different values of $\Omega$ when exogenous variation is driven by eqn. S15 in the main text. The approximation does a very good job for $\bar{a}$ near 1 and for longer cycle periods. Note that for longer cycle periods, the approximation for the variance of $a_t$ becomes better and there are longer autocorrelations at each time step, and so $c$ becomes more important.

### S3 Variance of the stochastic Ricker model

To approximate the variance in the stochastic Ricker model, we extend the approximation from section S2 "Variance of the exogenous cycle" to also include stochastic environmental variation. The stochastic Ricker model can be rewritten as

$$N_{t+1} = N_t \exp\left(A_t - bN_t\right) \tag{S31}$$

where $A_t = a_t + \sigma Z_t$ is now a random variable that includes stochastic variation in the environment as well as deterministic, exogenous variation. Remembering that $Z_t \overset{\text{i.i.d.}}{\sim} \mathcal{N}(0,1)$, $A_t$ is simply a translation of a normal and so is itself a normal such that, for any $t$,

$$A_t \sim \mathcal{N}\left(a_t, \sigma^2\right). \tag{S32}$$

To get an understanding of the variance of the stochastic Ricker model, we employ the same small variance approximation as in the previous section, but now where we have stochastic variation in addition to the deterministic, exogenous variation. As before, we re-scale population size so that we model relative density $N_t' = N_t/\bar{N}$, where $\bar{N} = \bar{a}/b$. As we show below, the long-term average of the sequence of $A_t$ converges on $\bar{a}$.

As part of the approximation, we will need the statistical properties of the long-term sequence of $\{A_t\}$, not just the statistical properties of any individual time point. Let $\bar{A}_T$ be the random variable giving the average of the sequence of random variables $A_t$ from $t = 1, 2, 3, \ldots, T$, i.e.,

$$\bar{A}_T = \frac{1}{T}\sum_{t=1}^{T} A_t. \tag{S33}$$

By the central limit theorem, $\bar{A}_T \to E[A_t] = \bar{a}$ in the limit as $T \to \infty$.

The other measure of the sequence we are interested in is $V_T^A$, the variance of the sequence from all time points up to $T$, i.e., $t = 1, 2, 3, \ldots, T$. Given our interest in long-time sequences, we calculate the variance of the sequence w.r.t. $\bar{a}$, the asymptotic value of the $\bar{A}_T$, which makes the math decidedly simpler than what is typical, which is using $\bar{A}_T$, the mean at that time $T$. However, this simplification has no practical effect on the conclusions. We define the variance of the sequence as

$$V_T^A = \frac{1}{T}\sum_{t=1}^{T}(A_t - \bar{a})^2. \tag{S34}$$

To determine the properties of the distribution of the sequence variance, first, standardize the difference of the square such that $V_T^A$ can be written as

$$V_T^A = \frac{\sigma^2}{T}\sum_{t=1}^{T}\left(\frac{A_t - \bar{a}}{\sigma}\right)^2, \tag{S35}$$

where $(A_t - \bar{a})/\sigma \sim \mathcal{N}\Big((a_t - \bar{a})/\sigma, 1\Big)$. We need only find the mean and the variance of $V_T^A$ to understand the long-time limit.

To do so, we need the mean and the variance of the sum

$$\sum_{t=1}^{T}\left(\frac{A_t - \bar{a}}{\sigma}\right)^2. \tag{S36}$$

The sum of squares of normal distributions with non-zero mean and unit variance is described a non-central chi-square distribution. Specifically, for $k$ normally distributed random variables, $U_i \sim \mathcal{N}(\mu_i, 1)$ $(i = 1, 2, \ldots k)$, the sum of their squares $Q_k = \sum_{i=1}^{k} U_i^2$ follows a non-central $\chi^2$ distribution with $k$ degrees of freedom and non-centrality parameter $\lambda = \sum_{i=1}^{n} \mu_i^2$. The mean of a non-central $\chi^2$ is $E[Q_k] = k + \lambda$ and the variance is $\mathrm{Var}(Q_k) = 2(k + 2\lambda)$. Using this fact, it follows that

$$\begin{aligned} V_T^A &= \frac{\sigma^2}{T} Q_T \\ Q_T &\sim \chi_T^2\left(\lambda = \sum_{t=1}^{T}(a_t - \bar{a})^2/\sigma^2\right). \end{aligned} \tag{S37}$$

Hence, the mean of $Q_T$ is

$$E[Q_T] = E\left[\sum_{t=1}^{T}\left(\frac{A_t - \bar{a}}{\sigma}\right)^2\right] = T + \sum_{i=1}^{T}\left(\frac{a_t - \bar{a}}{\sigma}\right)^2 \tag{S38}$$

Now we may use (S37) and (S38) to find the mean of $V_T^A$, which is

$$\begin{aligned} E[V_T^A] &= \frac{\sigma^2}{T} E[Q_T] \\ &= \frac{\sigma^2}{T}\left[T + \sum_{t=1}^{T}\left(\frac{a_t - \bar{a}}{\sigma}\right)^2\right] \\ &= \sigma^2 + \frac{1}{T}\sum_{t=1}^{T}(a_t - \bar{a})^2 . \end{aligned} \tag{S39}$$

Note that, by definition, $\lim_{T\to\infty} T^{-1}\sum_t (a_t - \bar{a})^2 \to \sigma_a^2$. Hence, it follows that $E[V_\infty^A] = \sigma^2 + \sigma_a^2$.

The variance of the sequence variance is given by

$$\mathrm{Var}(V_T^A) = \frac{\sigma^4}{T^2}\mathrm{Var}(Q_T). \tag{S40}$$

The variance of $Q_T$ is

$$\mathrm{Var}(Q_T) = 2\left(T + 2\sum_{t=1}^{T}\left(\frac{a_t - \bar{a}}{\sigma}\right)^2\right), \tag{S41}$$

which when substituted into (S40) yields

$$\begin{aligned} \mathrm{Var}(V_T^A) &= \frac{\sigma^4}{T^2}\left[2\left(T + 2\sum_{t=1}^{T}\frac{(a_t - \bar{a})^2}{\sigma^2}\right)\right] \\ &= \frac{1}{T}2\sigma^2\left(\sigma^2 + 2\frac{1}{T}\sum_{t=1}^{T}(a_t - \bar{a})^2\right). \end{aligned} \tag{S42}$$

Taking the limit as $T \to \infty$, it follows that $\mathrm{Var}(V_\infty^A) \to 0$. Therefore, the random sequence $V_T^A$ converges uniformly on its expected value, i.e.,

$$\lim_{T\to\infty} V_T^A \to \sigma_a^2 + \sigma^2, \tag{S43}$$

provided $\sigma_a^2 + \sigma^2 < \infty$, where $\sigma_a^2$ is the deterministic, exogenous component of the variability and $\sigma^2$ is the stochastic component.

We may now use the small variance approximation technique used in the section above for understanding the deterministic model with exogenous variation. There, we assumed small variation in $a_t$. Here, all is the same except now we have small variance in $A_t$ about its long-term sequence average, $\bar{a}$.

Employing the approximation from above, we have

$$\begin{aligned} \lim_{T\to\infty}\frac{1}{T}\sum_{t=1}^{T}(N_t' - 1)^2 = &\lim_{T\to\infty}\frac{1}{T}\sum_{t=0}^{T-1}(A_t - \bar{a})^2 + (1-\bar{a})^2\lim_{T\to\infty}\frac{1}{T}\sum_{t=0}^{T-1}(N_t' - 1)^2 \\ &+ 2(1-\bar{a})\lim_{T\to\infty}\frac{1}{T}\sum_{t=0}^{T-1}(A_t - \bar{a})(N_t' - 1) + o(\sigma_A^2). \end{aligned} \tag{S44}$$

Now recognize that the limit of the sum of squared deviation in $A_t$ over the sequence is $V_\infty^A \to \sigma_a^2 + \sigma^2$. Furthermore, provided that population density is bounded, the sequence $\{N'\}$ reaches a stationary distribution in the long-time limit such that the statistical properties of the set from $t = 0$ to $t = T - 1$ converges on the set $t = 1$ to $t = T$. Therefore, we may take the limiting variances of these sequences as identical. Last, recognize that the average product of deviations in $A_t$ and deviations in $N_t'$ are equal to the deviations in $a_t$ and $n_t'$. The reason they are identical is because the extra variation in the stochastic model is independent over time and independent of population size and therefore is statistically uncorrelated with population size. Using these facts, we can finalize the approximation as

$$\lim_{T\to\infty}\frac{1}{T}\sum_{t=1}^{T}(N_t' - 1)^2 = \frac{\sigma_a^2(1+c) + \sigma^2}{1 - (1-\bar{a})^2} + o\Big((A_T - \bar{a})^2\Big), \tag{S45}$$

where $c = \lim_{T\to\infty}\sum_{u=1}^{T} 2(1-\bar{a})^u \rho_a(u)$ is the constant describing the scaling effect of autocorrelation structure in the exogenous driver. Note that this approximation only applies when the sum $\sigma_a^2 + \sigma^2$ is small, and so works well for a smaller range of variation in both aspects of variability. Nonetheless, it predicts the qualitative nature of the effect of stochastic variation on population variation as illustrated by Figure S4.

### S4 Derivation population variance partition

To develop a variance partitioning scheme for an observed realization of a population trajectory, consider the deterministic map $n_{t+1} = F(n_t)$ in discrete-time. For the continuous time case, we can arrive at such a map by suitably discretizing time, and finding soluations to the continuous-time model (typically a differential equation) at the discretized time points. For this deterministic tractory, $n_t$, let $\bar{n}$ be the average population size.

Now let $N_{t+1} = F(N_t, Z_t)$ be a sample path of a stochastic version of the map with environmental variability given by the random variable $Z_t$. For this stochastic process, let $\bar{N}$ be the average population size. At each time point, we may rewrite the deviation in population density of the stochastic model from the mean $(N_t - \bar{N})$ as three components:

$$N_t - \bar{N} = \underbrace{(n_t - \bar{n})}_{i} + \underbrace{(N_t - n_t)}_{\text{ii}} + \underbrace{(\bar{n} - \bar{N})}_{iii}, \tag{S46}$$

which rewrites the deviation of the stochastic process from its mean as (i) the deviation of the deterministic process from its mean, (ii) the deviation of the stochastic process from the deterministic process at time $t$, and (iii) the deviation of the deterministic mean from the stochastic mean.

When population abundance is written as in eqn. (S46), the variance in population density can be found by squaring each side of the equation and taking the expectation of the result:

$$\begin{aligned} \mathrm{E}[(N_t - \bar{N})^2] =& \mathrm{E}\left[(n_t - \bar{n})^2\right] + \mathrm{E}\left[(N_t - n_t)^2\right] + \mathrm{E}\left[(\bar{n} - \bar{N})^2\right] + \\ & 2\mathrm{E}\left[(n_t - \bar{n})(N_t - n_t)\right] + 2\mathrm{E}\left[(n_t - \bar{n})(\bar{N} - \bar{n})\right] + 2\mathrm{E}\left[(N_t - n_t)(\bar{N} - \bar{n})\right]. \end{aligned}$$

For compactness and interpretability we prefer to rewrite the above equation as the following sum:

$$\mathrm{Var}(N) = \sigma^2_{\text{Det}} + \sigma^2_{\text{Stoc}} + \sigma^2_{\text{Nonlin}} + \sigma_{\text{Int}}, \tag{S47}$$

where

$$\begin{aligned} \mathrm{Var}(N) &= \mathrm{E}[(N_t - \bar{N})^2] \\ \sigma^2_{\text{Det}} &= \mathrm{E}\left[(n_t - \bar{n})^2\right] \\ \sigma^2_{\text{Stoch}} &= \mathrm{E}\left[(N_t - n_t)^2\right] \\ \sigma^2_{\text{Nonlin}} &= -(\bar{n} - \bar{N})^2 \\ \sigma_{\text{Int}} &= 2\mathrm{E}\left[(n_t - \bar{n})(N_t - n_t)\right]. \end{aligned}$$

- The term $\sigma^2_{\text{Det}}$ is the variance in the unobserved deterministic trajectory.
- The term $\sigma^2_{\text{Stoc}}$ is the variance in the observed stochastic trajectory, relative to the unobserved deterministic process at the same time point.
- The term $\sigma^2_{\text{Nonlin}}$ quantifies the effects of nonlinear dynamics. When the mean of the stochastic process is equal to the mean of the deterministic process, $\bar{n} - \bar{N} = 0$, then term is $\sigma^2_{\text{Nonlin}} = 0$. In general, the effects of nonlinear averaging will drive differences between $\bar{N}$ and $\bar{n}$.
- The final term, $\sigma_{\text{Int}}$ measures how much the the stochastic deviations $(N_t - \bar{n})$ covary with the deterministic deviations from the fixed point $(n_t - \bar{n})$. This term will be negative, i.e., exhibit variance damping, when stochastic fluctuations tend to be positive when the deterministic fluctuation is negative.

#### S4.1 Estimating variance components

We estimate the variance components in eqn. (S47) from the observed realization of a stochastic process, denoted as $N_t$, by first fitting a model to the data. The first challenge in estimating these components is defining the deterministic trajectory. Consider the Markov process $P(N_t|N_{t-1}) = g(N_{t-1};\, \Theta)\exp^{\varepsilon_t}$, where $\varepsilon_t \sim \text{Normal}(0, \sigma^2)$, and $\Theta$ is a vector of parameters associated with the expected value of the process. The deterministic trajectory is then the limit of $P(N_t|N_{t-1})$ as $\sigma$ goes to 0. The deterministic map is then $n_t = \lim_{\sigma\to 0} g(n_{t-1};\, \Theta)\exp^{\varepsilon_t} = g(n_{t-1};\, \Theta)$, where we have switched to lowercase variables for the population state to emphasize that it is no longer a random variable.

In practice the values of $\Theta$ are unknown so we use the estimated parameters, $\widehat{\Theta}$, to determine the trajectory of the process in the absence of stochasticity. In order to estimate the uncertainty in the variance components we draw from the sampling distribution of $\widehat{\Theta}$, and for each draw from the sampling distribution we calculate a deterministic trajectory, $\widehat{n}_t$, for all $t$. For each draw of a potential deterministic trajectory we calculate the variance components using the observed stochastic trajectory.

We consider two cases for the initial conditions of the deterministic trajectory in the manuscript. For the flour beetle dataset we measure the initial condition, $n_0$, exactly. The deterministic trajectory for $n_t$ is given by the $t^{\text{th}}$ iterative map $\hat{n}_t = g^t(n_0; \widehat{\Theta})$. In the case of the Lynx dataset, the initial conditions are observed with error. Therefore we draw the initial conditions from the sampling distribution, then iterate, $\hat{n}_t = g^t(\hat{n}_0; \widehat{\Theta})$.

We rescaled the estimated empirical variance components in eqn. (S47) by $\text{Var}(N_t)$. This rescaling allows us to interpret the components as the proportional contribution to the total variance. However, this rescaling is only well-defined when $\text{Var}(N_t) > 0$.

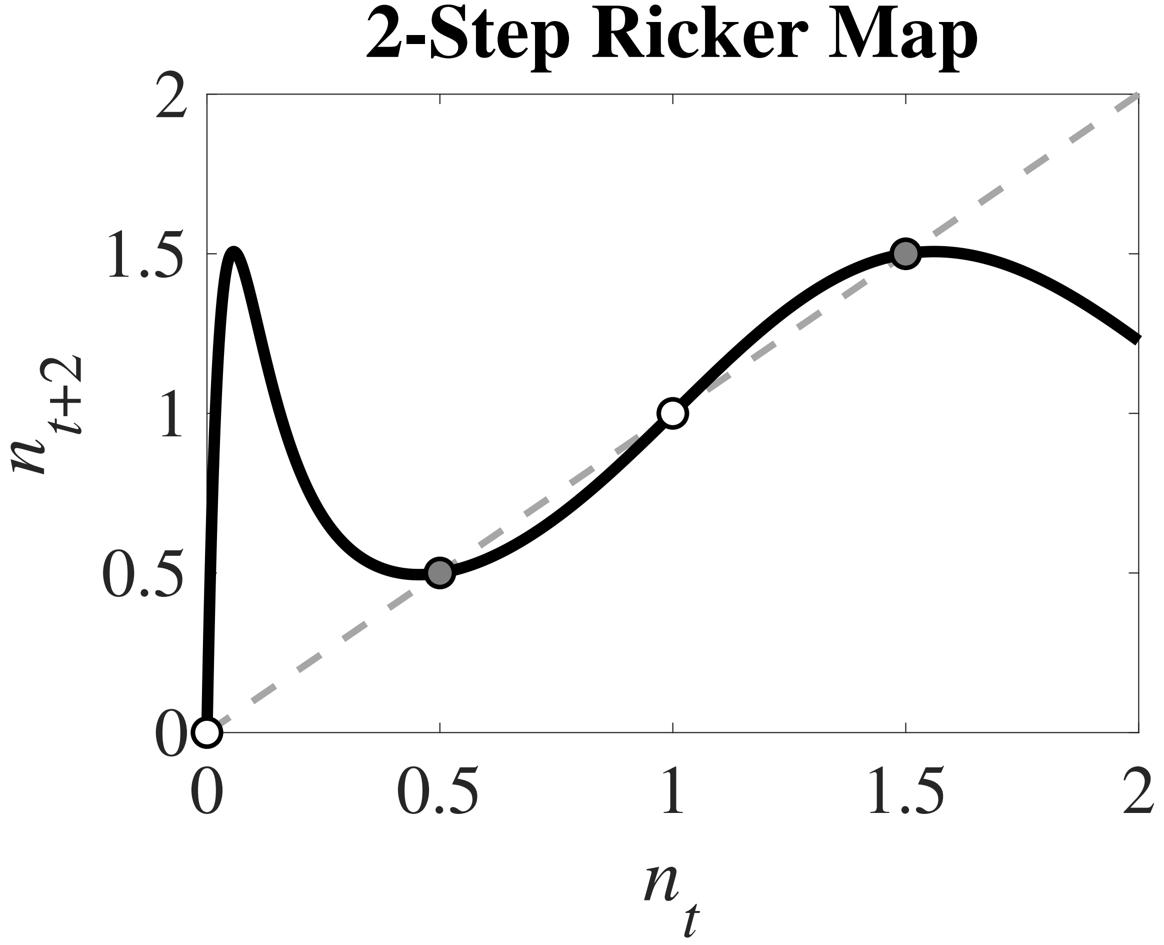

Figure S1: The two-step Ricker map in the region of parameter space giving rise to period-2 cycles. The two-step map is given by the black line and a reference 1:1 dashed line is provided to easily visualize equilibria. The two-step map has four equilibria, two stable (filled circles) and two unstable (open circles). Here $a = 2 \cdot \ln\{3\}$ and $b = a$, chosen so that the non-trivial unstable equilibrium is 1 and both stable equilibria are 0.5 units away from 1.

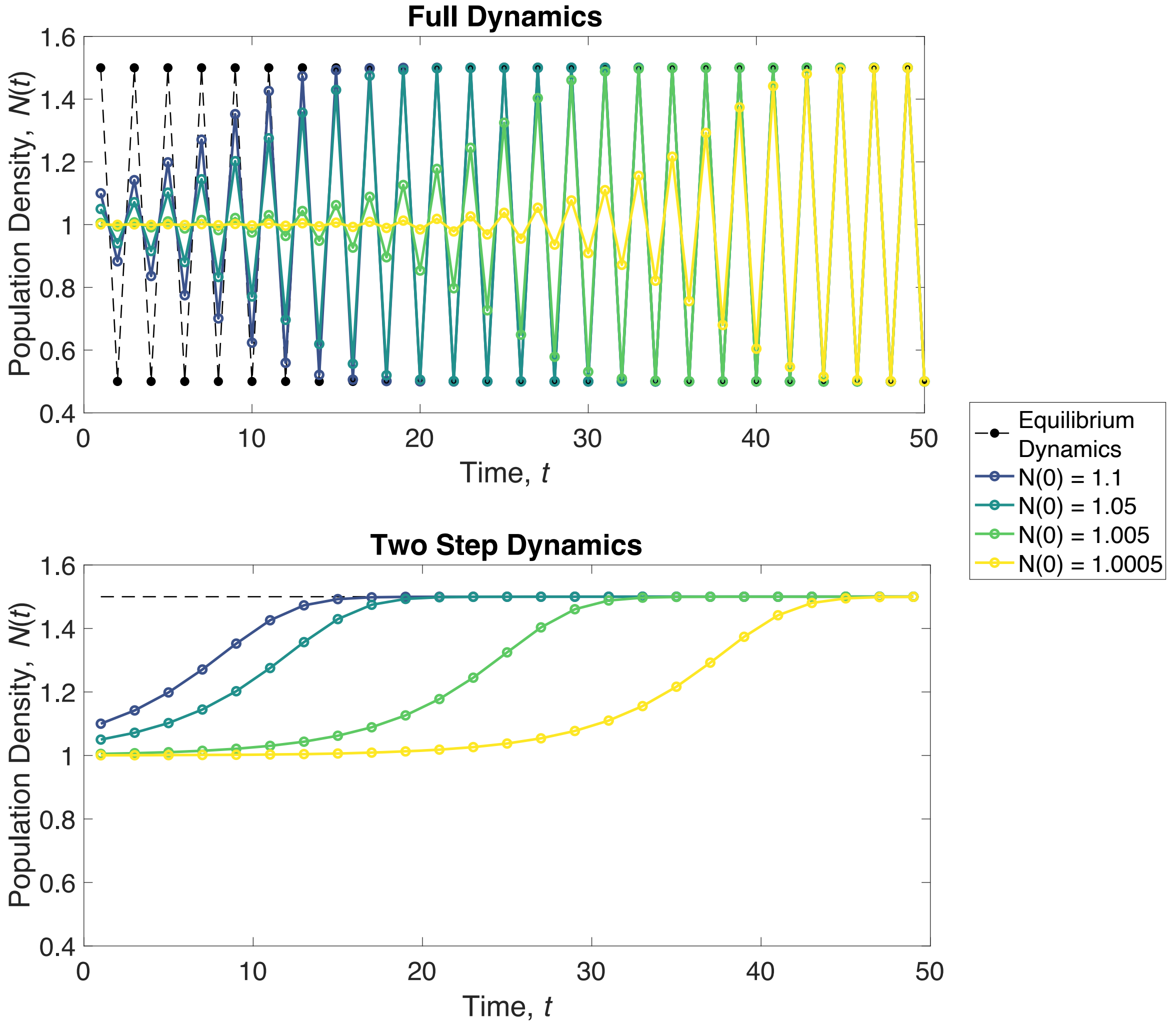

Figure S2: Transient dynamics of the Ricker model with a two-point cycle. Asymptotic dynamics shown by the dashed black line. Each line shows the transient trajectories towards the equilibrium for different initial values. Purpler colors are further away from the unstable equilibrium. Yellower colors are closer to the unstable equilibrium. The top panel shows the full dynamics. The bottom panel shows the two-step dynamics, which is population density every two time steps, which more clearly illustrates the transient nature of the dynamics. The unstable equilibrium is at $N = 1$. Only the odd values of $t$ are shown. Even values of $t$ show the same pattern but reflected across the y-axis along the value of 1. As in Figure S1, $a = b = 2 \cdot \ln\{3\}$.

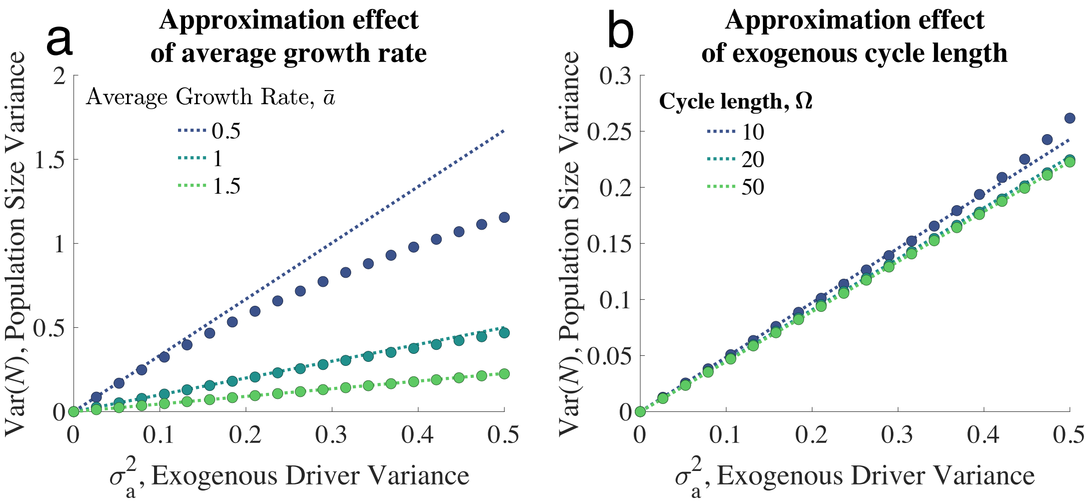

Figure S3: Correspondence between the variance in the time series of the Ricker model calculated from simulations (circles) and the analytical approximation (dotted lines; eqn. S30) under sine wave variation in $a_t$ (eqn. 3 in the main text). Panel a shows how well the approximation holds for different values of $\bar{a}$. Panel b shows how well the approximation holds for different cycle lengths, $\Omega$. Parameters: $\Omega = 20$ in panel a. $\bar{a} = 1.5$ in b. $T = 4000$ and $b = \bar{a}$ for all cases in both panels.

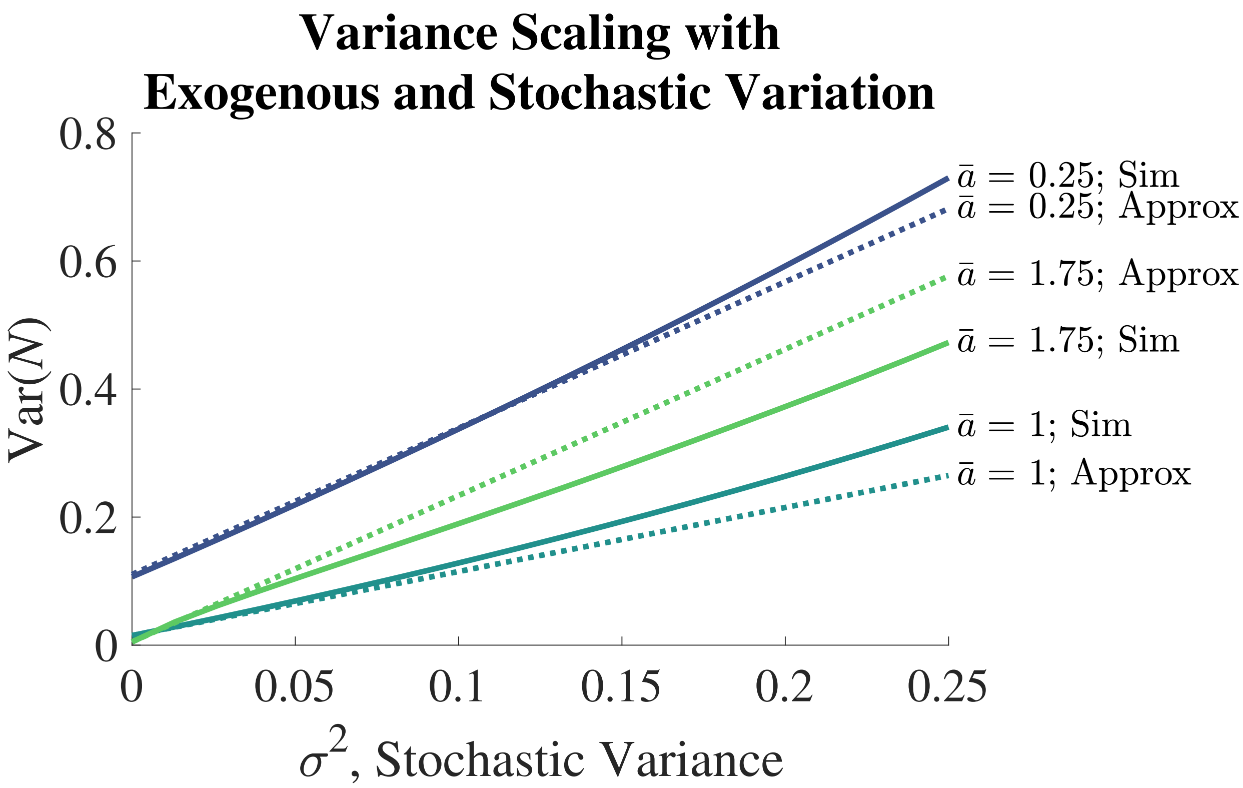

Figure S4: Scaling relationships for population variance under the model with sine wave exogenous variation. Variance from simulation of the model is given by solid lines. Approximation (S45) is given by dashed lines. The approximation does quite well explaining the qualitative relationship between stochastic variance, $\sigma^2$, and population variance, $\text{Var}(N)$. Other parameters: $\sigma_a^2 = 0.015$, $\Omega = 20$, and $b = \bar{a}$.
